# CATA: a comprehensive chromatin accessibility database for cancer

**DOI:** 10.1101/2020.05.16.099325

**Authors:** Jianyuan Zhou, Xuecang Li, Jiaxin Chen, Taisong Li, Weijie Zhan, Jianmei Zhao, Meng Li, Zhengmin Yu, Rui Yu, Haiying Zou, Wangkai Fang, Qiuyu Wang, Jianjun Xie

## Abstract

Chromatin accessibility is a crucial epigenetic concept that plays a biological role in oncology. As humans become more involved in cancer research, a comprehensive database is required to identify and annotate tumor chromatin accessible regions (CARs). Here, CATA was developed to provide cancer-related CAR annotation. Currently, CATA possesses 2,991,163 CARs, relevant clinical data, and transcription factor binding predictions for cancer CARs from 410 tumor samples of 24 cancer types. Furthermore, CARs were annotated by SNPs, risk SNPs, eQTLs, linkage disequilibrium SNPs, transcription factors, CNV, SNV, enhancer, and 450K methylation sites in our database. By combining all these resources, we believe that CATA will provide better service for researchers on oncology. Our database is accessible at http://bio.licpathway.net/cata/

## Introduction

Chromatin accessibility refers to the regions of a chromosome that have transcriptional activity[1], while these regions bind many transcription factors (TFs). Chromatin accessibility plays a very important role in tumorigenesis. It can help us discover more key information about epigenetics and cancer mechanisms. Researchers always used DNase-seq, to measure chromatin accessibility until that William J developed the Assay for Transposase-Accessible Chromatin using sequencing (ATAC-seq) technique in 2013[2]. Compared to DNase-seq, ATAC-seq has 4 major strengthens. First of all, the accuracy of ATAC-seq was consistent with that of DNase-seq but ATAC-seq experiment is easier to perform. Secondly, ATAC-seq kept simultaneous disclosure of accessible genomic location, DNA binding protein, transcriptional binding (TF) site interaction. Thirdly, ATAC-seq required fewer cell numbers. Fourthly, ATAC-seq showed great repeatability (R = 0.98) and also had good consistency with the DHS sequencing (R>0.79) [2].

ATAC-seq made it possible to assess the gene regulatory landscape in primary human cancers. Interestingly, chromatin accessibility changes are usually early cytological events in the context of various strain responses, stress responses, or developmental transitions [3]. In the early diagnosis and treatment of cancer, chromatin structure studies can provide valuable information [4]. However, the combinatorial effects of these phenomena in a specific biological function are poorly understood. Moreover, TFs are major players in the regulation of gene expression, and transcriptional regulation involves complex and detailed patterns of activity that bind to TFs. Numerous studies had indicated that TFs regulated genes by binding regulatory elements in the accessible genome [5]. There are a large number of regulatory elements, such as enhancers and promoters, located in ATAC-peak. Chromatin accessibility is diverse in different cancers, so there is a lot of transcriptional regulation information contained in CARs [6]. Identification of chromatin accessibility of cancer would provide ample insight into cancer mechanisms.

Several databases have stored chromatin accessibility data, such as Cistrome (Shenglin Mei et al., Nucleic Acids, 2017), TCGA(https://portal.gdc.cancer.gov/), and ENCODE (Carrie A Davis et al. NAR, 2018). They have become effective data sources for chromatin accessibility investigation. However, the existing resources did not focus on cancer and only contain limited cancer-related CARs, also they didn’t contain related comprehensive annotation information. Importantly, the cancer CAR usually contains a lot of regulatory elements [4,7] and could be bound by a lot of transcription factors, which might co-regulate target genes with the regulatory elements [8,9]. Here, we developed a comprehensive cancer chromatin accessibility database (CATA, http://bio.licpathway.net/cata/), which aims to provide a large number of available resources of cancer CARs and to annotate their potential roles in the regulation of a gene in cancer type-specific manner., CATA integrated data from 12 databases including TCGA, FUNTOM [10], 1000GENOMES [11], Jaspar [12] and Xena to annotated enhancers and TFs as well as mutation and methylation sites in CAR and developed CATA database. CATA contains 2,991,163 CAR and corresponding annotations for 410 tumor samples of 24 cancer types and supplies clinical data and survival analysis. CATA is also a comprehensive cancer ATAC-seq database that provides multiple functions, including storage, browsing, annotation, and analysis. It could be a powerful work platform for mining potential functions of CAR and explore relevant regular patterns about cancer.

## Materials and methods

### The collection of chromatin accessibility

We download chromatin accessible region data (bed file) from TCGA. First, The ATAC-seq data processing and alignment were performed using the PEPATAC pipeline (http://code.databio.org/PEPATAC/). The hg38 genome build used for alignment was obtained using Refgenie (https://github.com/databio/refgenie). Precisely, Bowtie2 was used to align to the hg38 human reference genome using “--very-sensitive - X 2000 --rg-id” options. Picard (http://broadinstitute.github.io/picard/) was then used to remove duplicates. Peak calling for 796 ATAC-seq profiles and 23 cancer types was performed to ensure high-quality fixed-width peaks with 501bp. For each sample, peak calling was performed on the Tn5-corrected single-base insertions using the MACS2 callpeak command with parameters “--shift −75 --extsize 150 --nomodel --call-summits --nolambda --keep-dup all -p 0.01”.The peak summits were then extended by 250 bp on either side to a final width of 501 bp, filtered by the ENCODE hg38 blacklist (https://www.encodeproject.org/annotations/ENCSR636HFF/), and filtered to remove peaks that extend beyond the ends of chromosomes. For overlapping peaks within a single sample, the most significant peak is kept and any peak that directly overlaps with that significant peak is removed. Then there is a set of fixed-width peaks for each sample. For each cancer, TCGA compiled a “cancer type-specific peak set” containing all of the reproducible peaks observed in an individual cancer type. For the overlapping peaks from different samples, TCGA kept the most significant peak. At last, using the same method, the ‘pan-cancer peak set’ was obtained from all cancer types that could then be used for cross-cancer comparisons.

### Annotation of chromatin accessibility regions

CARs were annotated both genetically and epigenetically through BEDTools [13], including common SNPs, risk SNPs, copy number variation, somatic mutation, motif changes, eQTLs, Transcription factors binding regions, methylation, enhancers, and CRISPR/Cas9 target sites, which prompted discovering potential functions of chromatin accessibility. Besides, interactive tables were utilized to present further detail.

### Enhancer

65,423 enhancers were collected from FANTOM5[10] and then converted to hg38 genome via LiftOver for the annotation.

### Transcriptionfactor-related data

Transcription factors binding regions were gained from FIMO [14] prediction. Besides, 5,797,266 TFBS [15] was downloaded from the UCSC[15] and convert to hg38 genome through LiftOver.

### CRISPR/Cas9 target sites

CRISPR/Cas9 target site [16], refers to RNA sequence within 200 bp of the genome region or DNA sequence in the transcription region, induction of which causes a precise cleavage of endogenous genomic loci. UCSC was employed to download CRISPR/Cas9 target sites, which were converted via LiftOver. Moreover, CRISPR tool was used in designation, evaluation, and clone for guidance sequence of the CRISPR/Cas9 system.

### Gene annotation

Three strategies were adopted to locate CAR associated genes. ROSE2 [17] gene-mapper method was applied in the prediction of associated genes including overlap, proximal, and closest.

### Common SNPs/Linkage disequilibrium SNPs/Risk SNPs

Total 38,063,729 common SNPs were downloaded from dbSNP [18] version 1.50. Gene linkage disequilibrium (LD) 1000 genome [11] project was calculated by 3 phased genotype information. Besides, minimum filtered SNP allele frequency of less than 0.05 (MAF) was accepted through VCFTools (v0.1.13). Moreover, plink (v1.9) was utilized to calculate SNPs with MAF > 0.05 in LD (r2 = 0.8) for 5 supergroups (Africa, ad mix America, East Asia, Europe, and South Asia). Risk SNP, genome-wide association studies (GWAS) resulting from GWAS catalog [19], and GWASdb [20] v2.0 set of planning were integrated into the table of human diseases/traits of SNP and insertion/deletion variant, which allowed functional annotation.

### Motif changes

Annotation of motif mutations was based on TRANSFAC [21] weight JASPAR [12] collection position weight matrix. The binding affinity of motif mutations, CAR located on the MAF> 1000 Genome Project 0.05 SNP stage, and 30-bp regions upstream and downstream of the SNP were calculated through R-package at-SNP. Finally, 254,545,586 motif changes were collected.

### TCGA series data

TCGA-related data including methylation data, RNA expression profile data, and Somatic-mutation-variation, Copy-number-variation, Clinical Information, ATAC-seq raw counts numbers, CAR about 24 cancer types were obtained from UCSC XENA (http://xena.ucsc.edu/). Additionally, similar data from involved in methylation and RNA expression profile was and other ATAC-raw count numbers were averaged based on cancer species and samples, respectively.

### EQTL

Human eQTL was downloaded and merged from GTEx [22] v5.0, HaploReg [23] for comparison in various tissues and PancanQTL [24] with TCGA eQTL gene relationship (https://tcga-data.nci.nih.gov/tcga). SNP eQTL was plotted and CAR was annotated which indicated an SNP gene regulated by the target gene as a potential associated CAR.

## Results

### A search interface for retrieving CAR

CATA offers 4 search methods to help users find CAR of cancer of interest. 1. Search for cancer CAR based on tumor samples (enter the cancer of interest) 2. Search for cancer CAR based on gene name (i. Enter the gene name of interest, gene symbol format, ii. Select the cancer type of interest). 3. Search for cancer CAR by transcription factors (i. Enter the transcription factor of interest. ii. Select the cancer type of interest.), 4. Search for relevant tumor CARs based on genomic coordinates (i. Enter the genomic coordinate information of interest, the CATA database supports the latest hg38 reference genome. ii. Select the cancer species). In the search results, CATA provides summative information about the CARs, including SNP, SNV, CNV, methylation, Motif, and so on. Click interest PEAK-ID can entering detail page. The Detail page consists of seven parts, including the CAR overview and annotation, RNA-expression, associative clinical data, survival analysis, methylation visualization, Upstream Transcription factor enrichment. 1. Accessible overview: CATA provides a preview of CAR, including raw counts number, presented by bar plots, and some summary information about the region. 2. Accessible region annotation: Provides chromatin in tabular form Information on the CAR, including SNP, motif, CNV, SNV, TFBS, etc., users can click the relevant knob to browse the corresponding information. 3. RNA-expression: We visualized the gene expression of the CAR by barplot. The user can select the gene annotated by ROSE2 to be divided into close, proximal, and overlap. 4. associative clinical data: We provide clinical information about patients with ATAC-seq, which is convenient for researchers to find potential cure targets. 5.survival analysis: CATA provides survival analysis of cancer CAR related genes to help researchers find genes with clinical value. 6.methation visualization: CATA uses barplot to visualize 450K methylation chip data, providing methylation in CAR, presented in 24 cancer forms. 7. Upstream Transcription factor enrichment: We used FIMO software to predict the binding factors of CAR and found some transcription factors and transcriptional cofactors that interact with transcription factors to form a visual network. Figure (contains: closet, proximal, overlap gene, combined motif, interact TF, and peak ID are marked by different colors).

### User’s friendly Explorations

CATA provides a user-friendly interface to help users to navigate quickly and easily. On the left side of the page, the user can distinguish between samples, via four options (Tissue type, Cancer type, Annotation, Chromosome), which can be clicked. The results are filtered and the user can click on the ‘Peak ID’ to jump to the details page of the CAR for more information.

### Personalized genome browser and data visualization

CATA provides the latest genome browsing area GIVE[25] to help visualize the open chromatin region. We provide 24 types of cancer and a total of 796 bigwig visualization files. Users can jump on the detail page, and view detailed regional information, or enter a region in the navigation bar and load the corresponding track for visualization. We grouped 23 cancers in detail and named the samples according to the TCGA-patient ID.

### Online analysis tools

CATA provides three analytical tools, including that: (1). Cell Pathway Analysis, in which the researchers analyzed pathways for binding transcription factors on the chromatin region and found upstream regulatory pathways. Users only need to input the of interest to enrich. We provide ten choices about pathway databases (KEGG, Reactome, NetPath, WikiPathways, PANTHER, PID, HumanCyc, CTD, SMPDB, and INO). According to the formula:

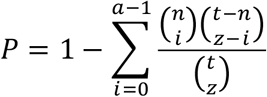

Where *t* is the number of genes of the entire genome, and *z* is the number of genes of interest, of which a gene involved in the pathway containing *n* gene. The calculated *P*-value is provided in the results page, along with the relevant genes, as well as the pathway ID (the pathway ID can be clicked into the view channel details); (2). Associated Accessible Region analysis, in which users can upload one or more genomic locations to our web. CATA also supported to upload a standard BED file. Then, CATA will provide brief information (CAR’s ID, genome location, and brief annotation information) that intersect with the user’s uploaded data. (3). Survival analysis, CATA allows users to select relevant genes for analysis in the detail page, which was implemented by built-in gepia2[26] (python3 package).

### Data download

Data from 24 types of cancer (ACC, BLCA, BRCA, CESC, CHOL, COAD, ESCA, GBM, HNSC, KIRC, KIRP, LGG, LIHC, LUAD, LUSC, MESO, PCPG, PRAD, SKCM, STAD, TGCT, THCA, UCEC, PAN) was available to download, including transcription factor information in CAR, and gene annotation information in CARs, users can click the icon to download interest data.

### System Design and Implementation

CATA was developed using MySQL 5.7.16 (http://www.mysql.com) and runs on a Linux-based Tomcat Web server (http://tomcat.apache.org/). We used JAVA 1.8 (https://www.java.com) for server-side scripting. The front end was designed and built using Bootstrap v4.1.1 (https://v4.bootcss.com). The network visualization was accomplished through Echarts[27] (https://www.echartsjs.com/) Chromatin interaction visualization was supplied by Genomic Interaction Visualization Engine[25] (GIVE) (https://zhong-lab-ucsd.github.io/GIVE_homepage/). We recommend using a modern web browser that supports the HTML5 standard such as Firefox, Google Chrome for the best display.

## Discussion

CATA is a tumor chromatin accessible regions database for cancer, storing 29,366,632 accessible chromatin regions from 24 types of cancers, including pan-cancer. Meanwhile, the CATA database annotates 2,936,663 CARs and stores the binding site of 1,678 transcription factors, which could be a good predictor for cancer-specific transcription factors. CATA also integrates data of methylation, SNP, SNV from TCGA and CATA has a favorable interactive interface for users. Four kinds of searching methods are provided by the database, which is based on cancer type, transcription factor, gene symbol, and advance search through genomic location, respectively. CATA also provides survival analysis of some CAR genes via the built-in gepia2 (python package). Also, CATA supports pathway analysis of transcription factors binding to CAR. All of that was aim to help cancer researchers easy to mine potential information for cancer mechanisms. However, there are still some deficiencies in CATA. Since the existing chip-seq data cannot perfect diverse for cancer species, It is believed that an explosion of these data will occur, or perhaps relevant chip-seq data and single-cell cancer ATAC-seq data will be added in the second edition to embellish the database, which enables better tumorigenesis mechanism and cancer markers mining.

**Figure 1.**
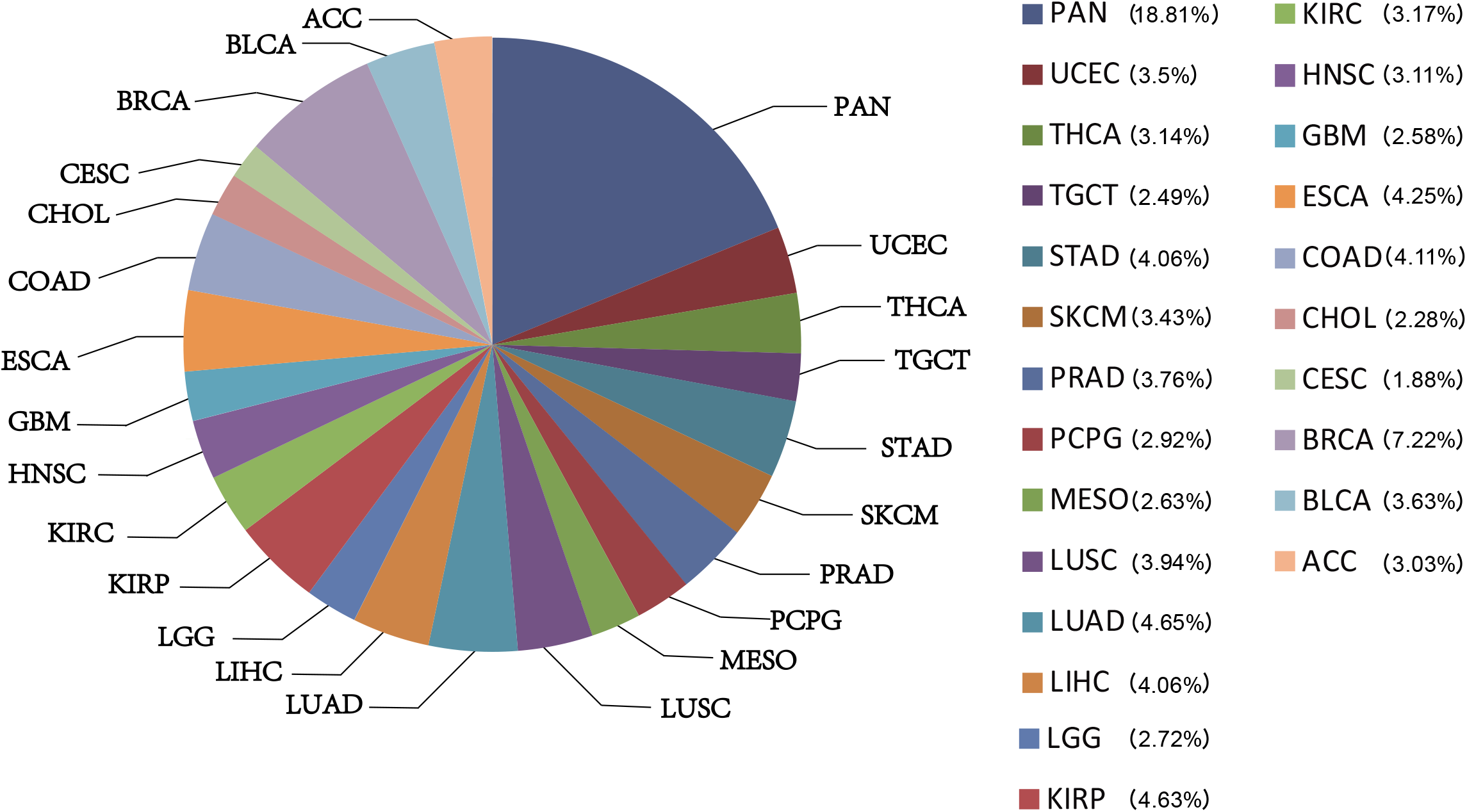
The percentage of chromatin accessible region for per cancer type.

**Figure 2.**
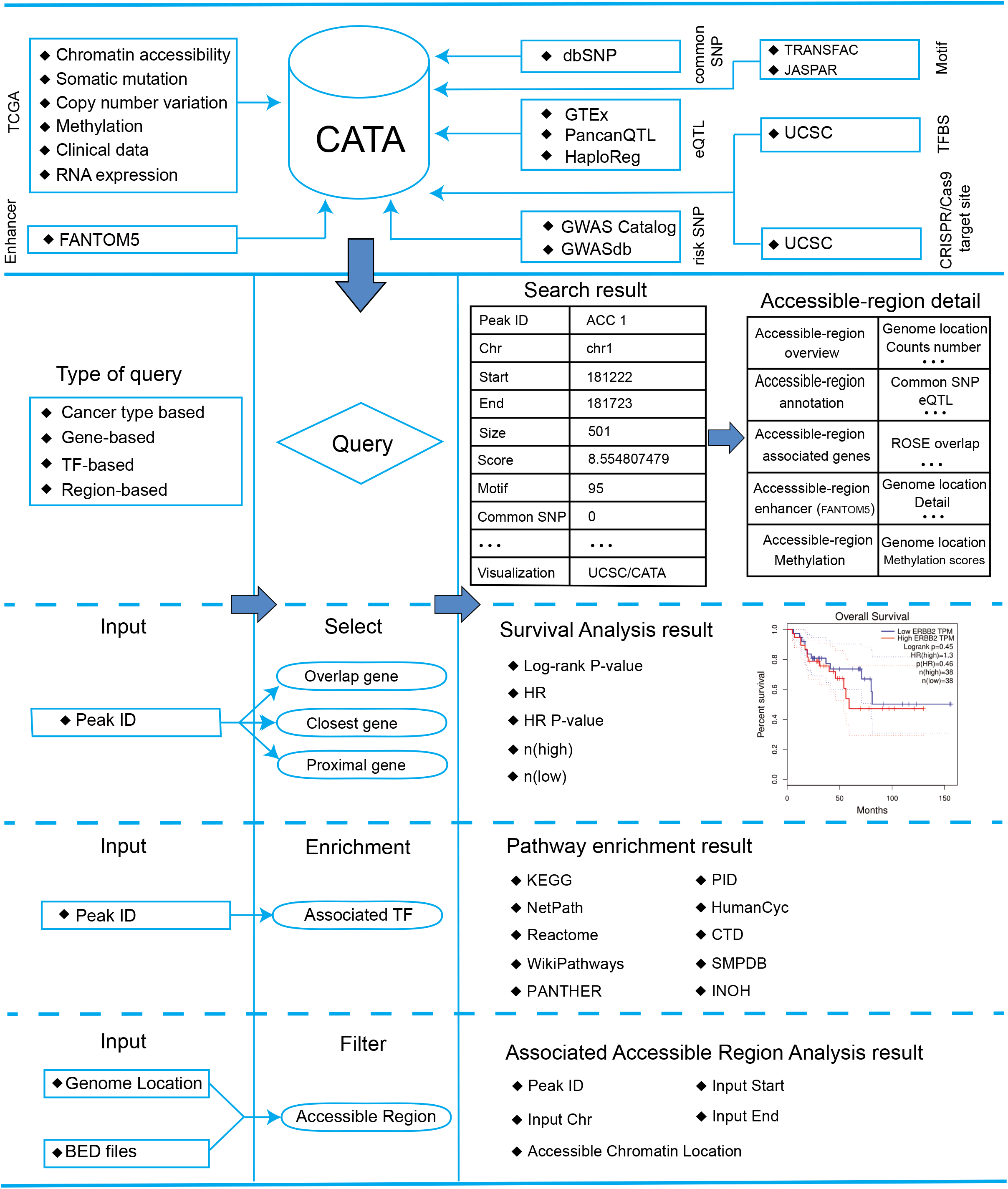
Database content and construction. CATA provide chromatin accessible region of cancer based on TCGA ATAC-seq data, Genetic and epigenetic annotations of accessible regions were collected or calculated including common SNPs, eQTLs, risk SNPs, LD SNPs, TFs, CNV, SNV, methylation sites, Enhancer. CATA also provide ATAC-seq samples associated clinical data. CATA integrates multiple functions including storage, search, download, statistics, visualization, browse and analysis.

**Figure 3.**
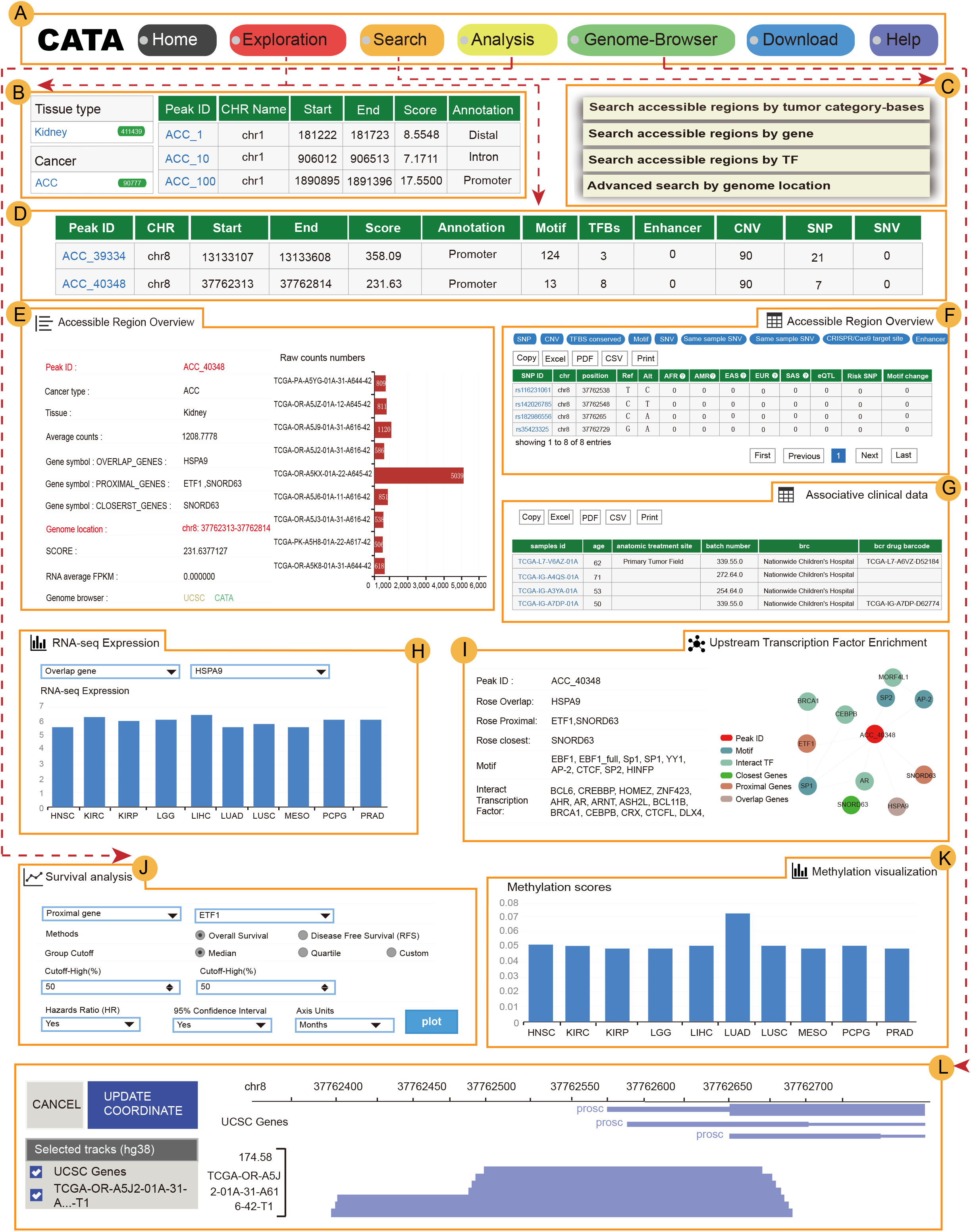
The main function and usage of CATA. (A) The navigation bar of CATA. (B) CATA provide user’s friendly Explorations. (C) Users query using five methods: ‘Search by Cancer type’, ‘Search by GENE’, ‘Search by TF’, ‘and ‘Advanced search by genome location’. (D) The search results. (E) Overview of chromatin accessible region. (F) Interactive table of chromatin accessible region, related annotation information. (G) The visualization of RNA-expression.(H) The table of clinical data. (I) Associated gene survival analysis. (J) The visualization of Methylation level. (K) Upstream Transcription factors enrichment. (L) Data download page. (M) Personalized genome browser-GIVE

## Acknowledgments

We thank TCGA for sharing their cancer chromatin accessibility data.

We thank Richard A. Young and his colleagues for sharing ROSE program with this work.

We thank Xiaoyi Cao help us to deploy Give genome-browsers.

We thank Zemin Zhang and his colleagues for sharing GIPIA2(python package) to this work.

## Key points

- CATA is the first comprehensive resource for chromatin accessibility for cancer.
- CATA provides TF-binding sites, methylation levels, enhancers, CNV, SNV, and SNP for CAR.
- CATA documents 2,991,163 CARs from 410 cancer samples.
- CATA genome-browser provides visualization of chromatin accessibility for 23 types of cancer.

## Funding

This work was supported by grants from the National Natural Science Foundation of China (81871921); the Natural Science Foundation of Guangdong Province-Outstanding Youth Project (2019B151502059) and the Basic & Applied Basic Research Programs of Guangdong province (2018KZDXM033).

## Notes

### Competing Interest Statement

The authors have declared no competing interest.

